# Low frequency somatic copy number alterations in normal human lymphocytes revealed by large scale single-cell whole genome profiling

**DOI:** 10.1101/2021.11.10.468149

**Authors:** Lu Liu, He Chen, Cheng Sun, Jianyun Zhang, Juncheng Wang, Meijie Du, Jie Li, Lin Di, Jie Shen, Shuang Geng, Yuhong Pang, Yingying Luo, Chen Wu, Yusi Fu, Zhe Zheng, Jianbin Wang, Yanyi Huang

**Author notes:** Correspondence to (Y.F.), (Z.Z.), (J.W.), (Y.H.). These authors contributed equally to this work (Lu Liu, He Chen, Cheng Sun, Jianyun Zhang).

## Abstract

Genomic-scale somatic copy number alterations in healthy humans are difficult to investigate because of low occurrence rates and the structural variations’ stochastic natures. Using a Tn5-transposase assisted single-cell whole genome sequencing method, we sequenced over 20,000 single lymphocytes from 16 individuals. Then, with the scale increased to a few thousand single cells per individual, we found that about 7.5% of the cells had large-size copy number alterations. Trisomy 21 was the most prevalent aneuploid event among all autosomal copy number alterations, while monosomy X occurred most frequently in over-30-year-old females. In the monosomy X single cells from individuals with phased genomes and identified X-inactivation ratios in bulk, the inactive X Chromosomes were lost more often than were the active ones.

## Introduction

Genomic alterations, including copy number alterations (CNAs) and point mutations, are the major drivers of many cellular malfunctions (Conrad et al. 2010; Sudmant et al. 2015). Tumor cells, for example, usually carry many CNAs and point mutations (Beroukhim et al. 2010; Waddell et al. 2015), many of which are oncogenic. After decades of study, researchers now recognize that point mutations accumulate in normal cells through polymerase replication errors and damaged DNA. Many point mutations barely affect cells, while others, located at critical locations, can transform cells (The Wellcome Trust Case Control Consortium 2010; Klopocki et al. 2011). The scenario for CNAs in normal cells is less clear. While large CNAs are extremely rare in humans, thus suggesting their destructive potential in cells (Zhang et al. 2009; Girirajan et al. 2011; Zarrei et al. 2015), their occurrences in normal cells may have been underestimated due to technical constraints.

Recent advances in single-cell sequencing have greatly extended our understanding of cellular complexity (Vitak et al. 2017; Zahn et al. 2017; Chen et al. 2017). However, few studies have reported the heterogeneities of nuclear genomic variations in single cells (Rohrback et al. 2018; Laks et al. 2019). Various technologies have estimated that the frequencies of somatic CNAs in the human brain vary between 2% and 40% (McConnell et al. 2013; Cai et al. 2014; Knouse et al. 2014; van den Bos et al. 2016; Chronister et al. 2019). The current lack of a scalable and robust method to perform uniform single-cell whole genome amplification (WGA) is the main challenge to improving the accuracy and precision of somatic CNA identification.

In this report, we present a high-throughput single-cell WGA and sequencing method: Tn5-transposase–assisted single-cell whole genome sequencing (Tasc-WGS). We demonstrate the power of Tasc-WGS by showing the results of a large scale investigation that identified rare CNA events in the lymphocytes from 16 cancer-free individuals of different ages and sexes. We portrayed the CNA pattern of normal lymphocytes and discovered hot spot regions. Combined with haplotype and transcriptome data, we were able to reveal the biological bias of aneuploid events in Chromosome X.

## Results

### A high-throughput pipeline for single-cell somatic CNA analysis

We optimized our Tasc-WGS method by forgoing both pre-amplification and library quantification, and by directly tagmenting the double-stranded genomic DNA of single cells, thus greatly simplifying the experimental process and increasing throughput (**Fig. 1A**). Previous studies had shown that the library construction protocol performed in a microfluidics system enables uniform amplification of a single-cell genome (Zahn et al. 2017; Laks et al. 2019). We expanded that protocol to 96-well plates, and our subsequent performance evaluation verified that our protocol provided even amplification and little contaminated data, as well as a marked increase in throughput that needed no special instrumentation (**Supplementary Fig. S1A, B, C**). We typically processed about 2,000 single cells in a single experimental run and used cellular barcodes to differentiate each cell’s sequence reads (**Table S2**).

**Fig. 1.**
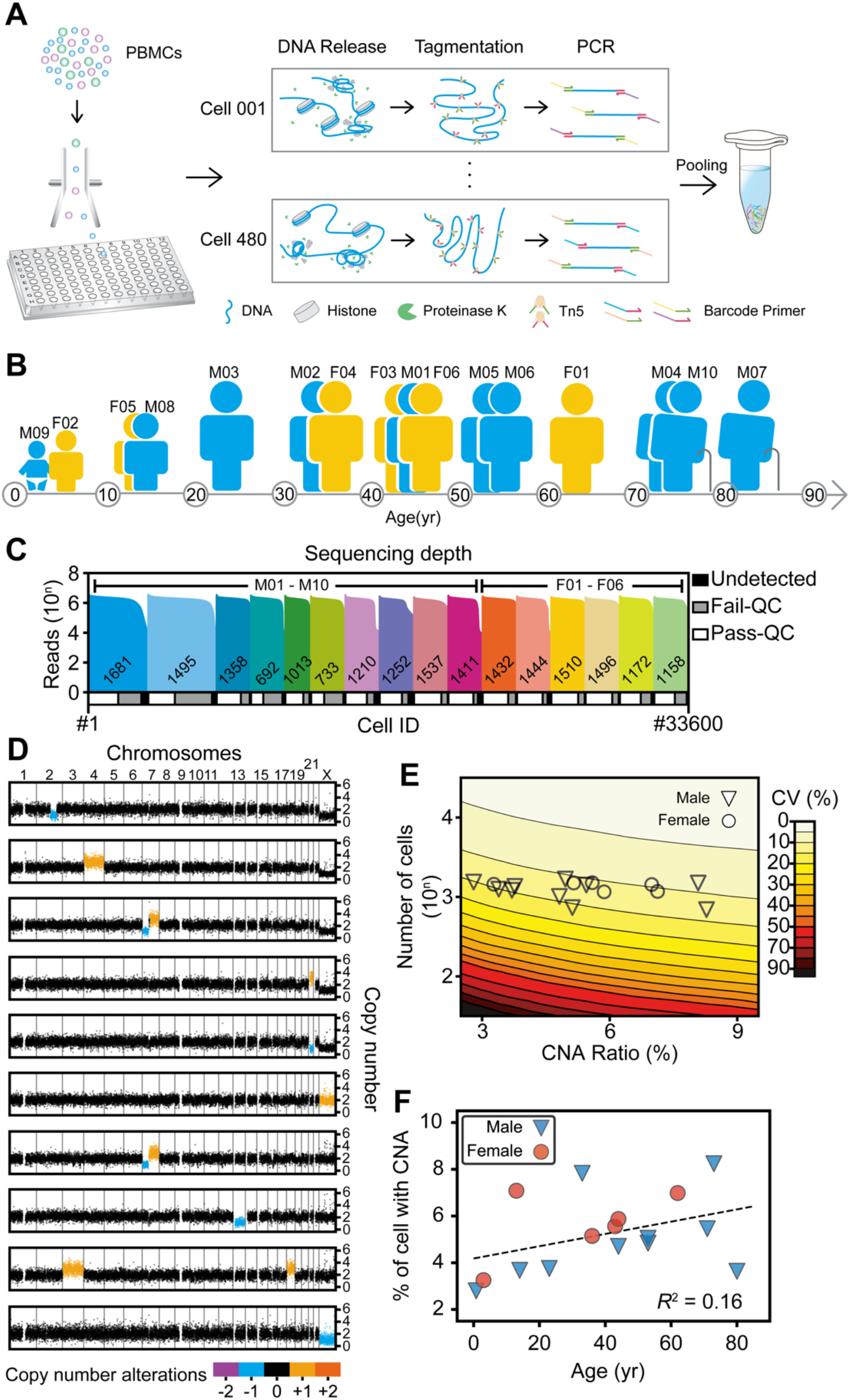
Overview of the study design. **(A)** Experimental flow used in this study. Lymphocytes were sorted to 96-well plates using fluorescence-activated cell sorting to obtain 1 cell per well. Thousands of those single-cells were lysed, tagmented, barcoded, and amplified in their wells and then pooled for second-generation sequencing. PBMC, peripheral blood mononuclear cells. **(B)** Cartoon showing the blood donors’ ages, sex, and ID numbers: 6 females (F) and 10 males (M) between 9 months and 80 years of age. **(C)** Sequencing depths and cell numbers for all samples in this study. The top histogram shows the reads counts distribution and the diagram underneath represents the relative proportions of cells after filtering. Undetected, low read counts; Fail-QC, failed quality filtering; Pass-QC, passed quality filtering. **(D)** Copy number profiles of representative cells with copy number alterations (CNAs) in colors. **(E)** Coefficients of variation (CV) of CNA ratio estimations. The contour plot shows both the theoretical CVs of CNA ratios (calculated by simulation) and the sample sizes (number of cells). The symbols show the real CNA ratios and sample sizes for each sample in this study. The large sample size in our study ensured that the CV was in a relatively small interval, thus providing acceptably accurate estimations. **(F)** Ages and autosomal CNA percentages of each sample. The dashed line indicates a weak linear relationship between age and CNA ratios.

With a shallow sequencing depth (~ 0.1×; i.e., 0.3 Gb per cell), we obtained an average 3.50% ± 1.50% (CI = 95%) genome coverage and detailed copy number profile with 200-kb bins. The coverage was uniform across the whole genome (**Supplementary Fig. S1A**), with no crosstalk between samples (**Supplementary Fig. S1B, Supplementary Notes**). We combined circular binary segmentation and hidden Markov model algorithms to further reduce false identification of unambiguous copy number losses or gains (**Supplementary Fig. S1D, E, Supplementary Notes**). Most of the quality filtering of single cell sequencing data was done with combinatorial criteria, including mean absolute deviation of pairwise difference, degree of ploidy abnormality, and degree of fragmentation (**Supplementary Fig. S1F, G, Supplementary Notes**).

Using a primary tumor sample cell-line and split-cell genome DNA as controls (**Supplementary Fig. S2**), we further validated the robustness of our Tasc-WGS protocol. First, we generated 96 libraries of 6-pg bulk, dilute, blood gDNA and then used it to show that tagmentation reactions and PCR occurred uniformly across the whole genome (**Supplementary Fig. S2B, C**). To evaluate the sensitivity of Tasc-WGS, we sequenced both a bulk and 29 single-cell samples of the HEK293 cell line, subsequently finding our method capable of detecting various types of CNAs ranging in size from 1 Mb to 26 Mb: CN gains ≥ 3.0, range 3.06–7.91, median = 3.60; copy number losses ≤ 1.3, range 1.20–1.30, median = 1.28 (**Supplementary Fig. S2D, E**). To evaluate the specificity of Tasc-WGS, we conducted CNA calling on a GM12878 B-lymphoblastoid cell line. First, bulk DNA genome profiling of GM12878 proved the diploid karyotype, and then we found some subpopulations with shared CNAs, as well as some unique CNAs, in the 49 GM12878 single-cell sequences (**Supplementary Fig. S2F**). Most of those unique CNAs ranged in size from 0.6 Mb to 10 Mb (**Supplementary Fig. S2G, Table S4**). So, to examine whether those small CNAs could have been data point noise mistakenly called by the algorithm, we simulated normally distributed copy number profiles that had multiple noise levels (Nilsen et al. 2012, **Supplementary Fig. S3**). As expected, noisier data tended to have higher false positive (FP) rates and FP CNAs were all small, ranging from 0.6 Mb to 4.8 Mb (mean = 2.3 Mb) (**Table S5**). Although it is possible that small CNAs (<10 Mb) occurred more because they affected fewer genes and thus affected cell survival less than larger CNAs do, it is hard to distinguish between true- and false-positive calls of small CNAs. Therefore, for the lymphocyte samples, we decided to include only CNA calls larger than 2 Mb.

### Somatic CNA events occurred commonly in lymphocytes

We examined 33,600 single lymphocytes sorted from the blood of 16 cancer-free individuals (10 males and 6 females; 1,440–3,840 single lymphocytes per individual) aged 9 months to 80 years (**Fig. 1B, Table S1**). Among them, 31,125 cells (92.6%) had more than 0.3 million reads aligned to the reference genome, and 20,594 cells (61.3%) passed the aforementioned quality filtering criteria for CNA analyses (**Fig. 1C, Supplementary Fig. S4A–E, Supplementary Notes**).

We found somatic CNA-containing lymphocytes in all individuals and identified 4,809 cells (24.0% on average, 12.9%–44.5% for different individuals) harboring small CNAs (2–10 Mb) and 1,500 cells (7.3% on average, 3.3%–15.2% for different individuals) harboring large CNAs (>10 Mb) (**Fig. 1D, Supplementary Fig. S4F**). Furthermore, we observed a few cells, from different individuals, that carried similar CNAs. Some other cells carried multiple CNAs across the whole genome (**Fig. 1D**). As in previous reports (Knouse et al. 2016), copy number deletions occurred much more often than did copy number amplifications (**Supplementary Fig. S4G, H**), and the ratios of CNA-containing cells were similar between males and females (**Supplementary Fig. S4I**). Because large CNAs may affect more genes and cause more serious problems than small CNAs, we focused on the large somatic CNAs (excluding Chromosome Y due to technical challenges) that we had identified in 1,397 lymphocytes.

We examined a large number of cells to minimize sampling error and to accurately assess the ratio of CNA-containing lymphocytes. To that end, we ran a simulation to determine the optimal number of cells that had to be sequenced to accurately assess the ratio of CNA-containing cells, and found that a smaller sample invariably yields uncertain assessment results (**Fig. 1E**). A throughput of about 1,000 or more cells per sample was ideal to achieve a coefficient of variation (CV) below 20%, and thus accurate CNA assessment with occurrence ratios less than 10%. We then checked whether those somatic, megabase-size CNAs in lymphocytes were age related (Machiela et al. 2015; Vattathil et al. 2016), and found a relatively weak correlation (**Fig. 1F**). All our observations suggested that CNAs were common in lymphocytes of cancer-free individuals.

### Large-size autosomal somatic CNAs occurred randomly in lymphocytes

We further analyzed CNA profile similarities between cells to try to capture clonal amplification signatures. Dimension-reduction analysis of single-cell copy number profiles produced a few clusters based on large-CNA patterns (**Fig. 2A, Supplementary Fig. S5A, Supplementary Notes**). Chromosome 21 and Chr X CNAs separated from the others, mostly because of their distinct aneuploidy patterns, but we observed no other obvious clustering based on sex, sample, or chromosome (**Supplementary Fig. S5A**). Using pairwise Euclidean distances of CNA profiles, we found 51 cell clusters, all with potential clonal cell amplifications (**Fig. 2B–D, Supplementary Fig. S5B, Supplementary Notes**). Although clonal amplifications among lymphocytes are not common (cell ratio = 2.3%, median clone size = 4), we found that most individuals (8/10 males and 6/6 females) exhibited such events, with clone sizes ranging from 3 to 105 cells. Aside from the characteristic aneuploid clones of Chr 21- and Chr X-containing cells, the largest clone, in F01’s Chromosome 6, contained 11 cells with an approximate 35 Mb loss (**Fig. 2E**).

**Fig. 2.**
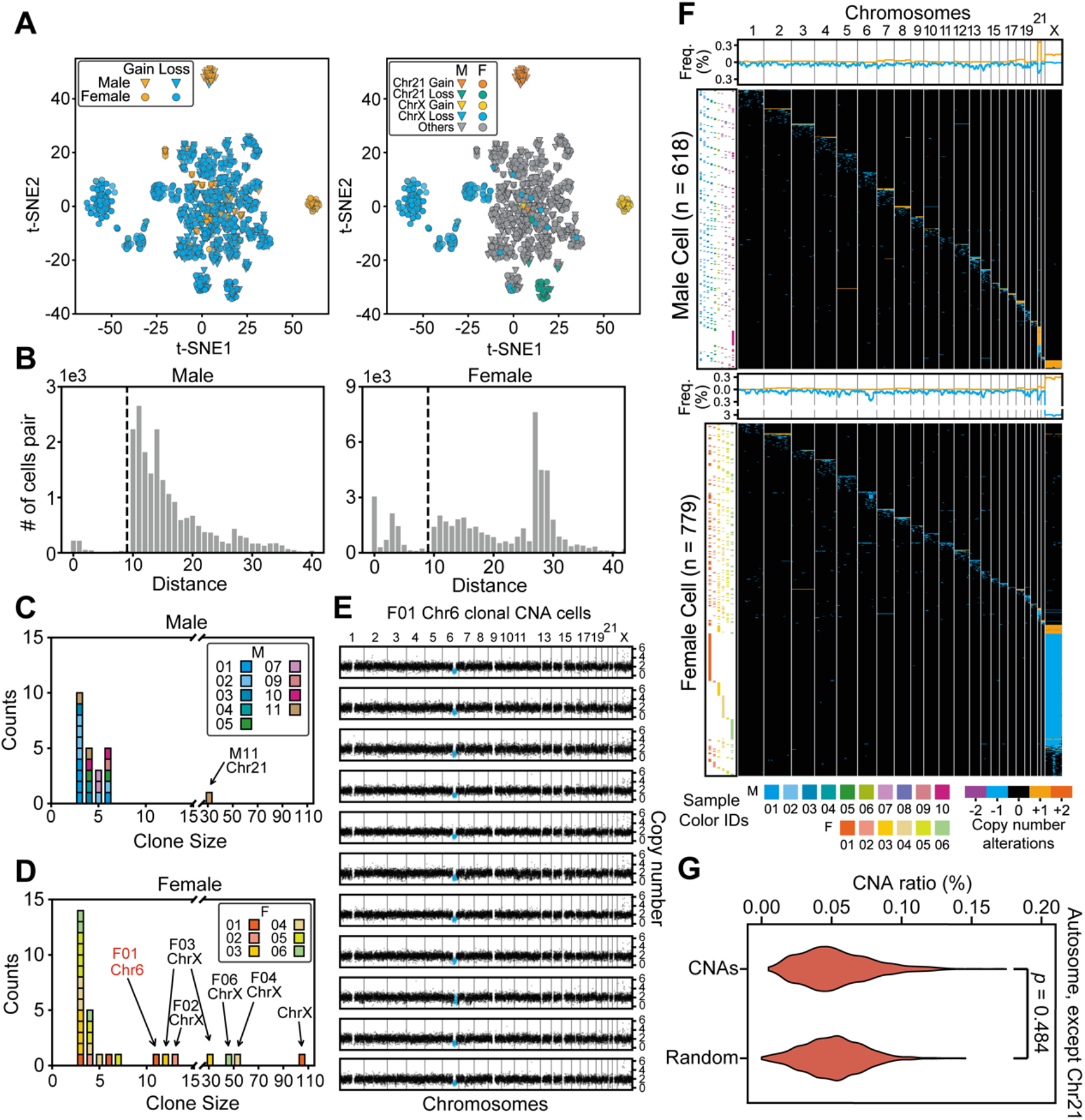
The landscape of cells with copy number alterations (CNAs). **(A)** Low-dimensional representations produced through multidimensional scaling of the copy number profiles for cells with >10-Mb CNAs. Colors label different types and locations of CNAs. t-SNE, *t*-distributed stochastic neighbor embedding. **(B)** The distributions of Euclidean distances between cell pairs. Two clusters were clearly separated by distance (*d*) < 10 (indicated by dashed lines) in both male and female samples. **(C, D)** The sizes (number of cells in a clone) and counts of clonal CNAs. Each block in the bar plot represents a clone. Most clonal CNAs with bigger clone sizes were on chr 21 and chr X, although there are some small clones with cell numbers of about 3–5. **(E)** Copy number profiles of each clonal CNA on chr 6 in 11 cells from F01. Each graph represents 1 cell and the blue dots represent the regions with copy number alterations. **(F)** Overview showing every cell with >10-Mb CNAs each. The heatmap (in black) demonstrates the genome patterns for cells with CNAs, which are labeled by different colors for gain and loss. Cells were sorted according to the chromosome fraction carrying CNAs. Each row represents one cell. The scatter plot (left) shows the cells’ sample IDs. The density curve along the top of each heatmap shows the aggregate frequency, at 200-kb bins, among all samples for each genomic locus. **(G)** CNA frequency distribution for each genomic locus in all autosomes except for Chr 21. The distributions of this study’s CNAs and of randomly generated CNAs were not significantly different (Mann–Whitney *U* test), thus indicating a random generation mechanism.

Somatic CNA events were scattered across every chromosome (**Fig. 2F**), although all CNAs together could cover almost the entire genome (99.4%). For each genomic locus that contained CNA events, the occurrence frequency was less than 1% (0.0%– 1.0%), except for Chr X (1.3%–1.8%) (**Supplementary Fig. S5C**). We detected no shared CNA hotspots among the participants, and CNA distributions were no different than a random distribution (**Fig. 2G**), except for the obviously higher frequencies at Chrs 21 and X. In keeping with the random generation mechanism, longer chromosomes contained proportionally more CNAs than shorter ones did (in a linear relationship, **Fig. 3A**), while Chr 21 and Chr X exhibited significantly higher occurrence rates than other chromosomes. We then examined whether certain chromosomes were more prone to CNAs than others and found that, except for Chr X and Chr 21, all other autosomes showed similar CNA count densities (**Fig. 3B, Supplementary Fig. S5D**).

**Fig. 3.**
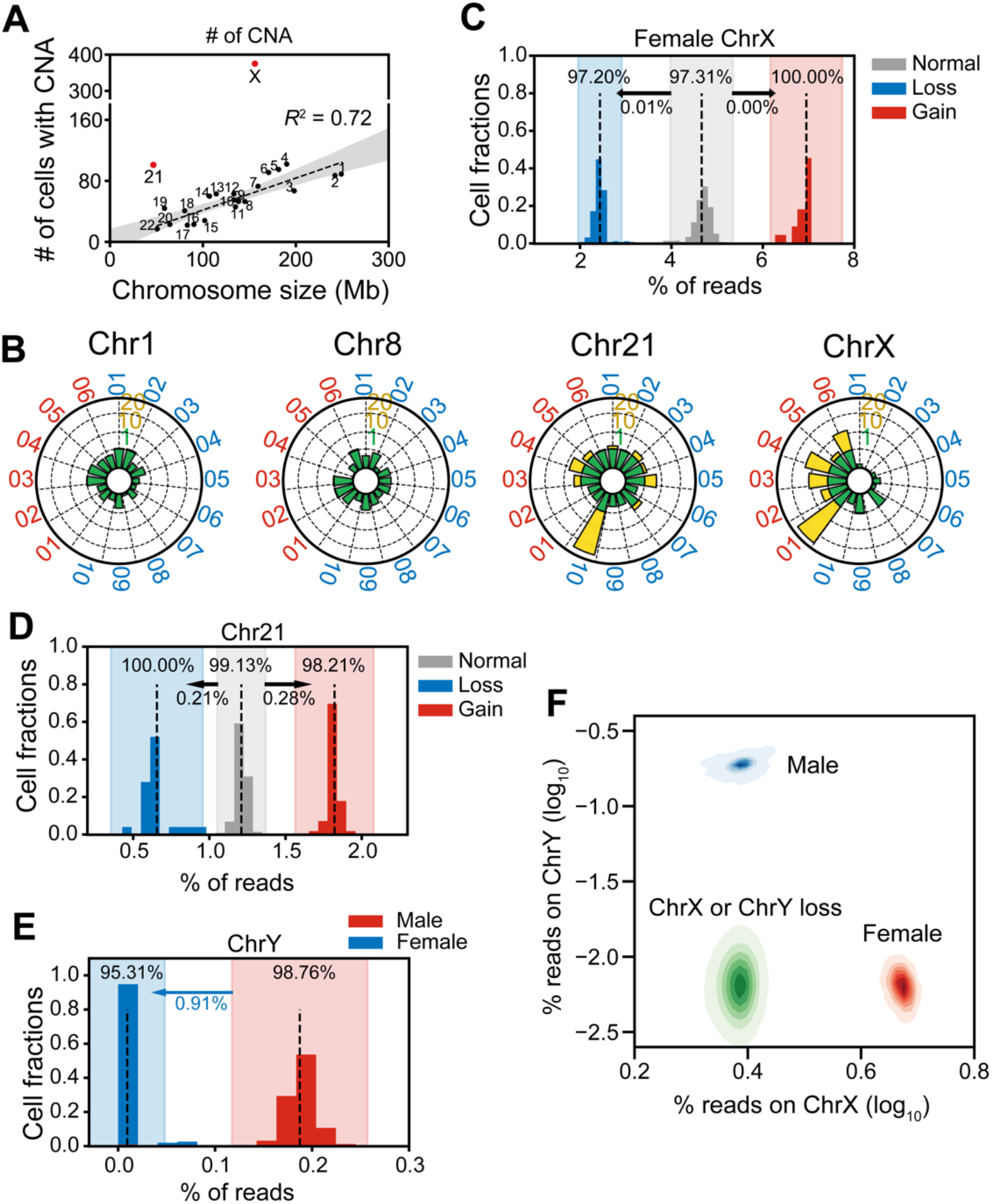
Clonal analysis of cells with CNAs. **(A)** Numbers of cells with CNAs for each chromosome. The dashed line indicates a significant correlation between chromosome size and numbers of CNAs. Chromosomes 21 and X (red) had markedly higher numbers of CNAs than the rest of the chromosomes had. **(B)** Radar plots show normalized CNA counts in 4 chromosomes from every individual labeled around each plot (blue, male; red, female). Internal colors denote a graded scale of CNA counts (green, 0–1; yellow, 1–20). **(C–E)** Reads counts distributions on Chromosomes X **(C)**, 21 **(D)**, and Y **(E)**. Each histogram shows the normalized reads distributions for each type of cells (gain, loss, and normal). Dashed lines indicate the means of each type of cell. The color shade indicates the confidence interval within 3 standard deviations. The percentage in each shaded bar indicates the fraction of cells identified as normal/loss/gain using reads counts. The percentages under the arrows shows the fractions of normal cells found, using reads counts, in the loss/gain confidence intervals. For Chromosome Y, those male cells that fell into the female confidence interval were identified as Chromosome Y losses. **(F)** Reads densities of Chr X and Chr Y of normal male cells (blue), normal female cells (red), and abnormal male and female cells with Chr Y or Chr X loss, respectively (both green).

CNA size distribution showed that copy number amplifications affect larger CNAs more than do deletions and they were usually aneuploidies (**Supplementary Fig. S6A**). Specifically, excepting Chr 21, 57.6% (38/66) of the autosomal copy number amplifications were aneuploidies, while only 2.0% of the deletions were (22/1123) (**Supplementary Fig. S6B, C**). These results suggest that the mechanism for copy number gain may be different than that for loss.

### Aneuploidy occurred mainly at Chr 21 and the sex chromosomes

We identified somatic aneuploidy in almost all chromosomes, but with a 2.4% (n = 498) cellular occurrence rate, it can be easily missed if the experimental throughput is not large enough. We verified low-frequency autosomal aneuploidy using fluorescence in situ hybridization assays, and quantitatively confirmed the copy number gains and losses in Chr 3, 8, 13, 18, 21 and X (**Supplementary Fig. S5E, Table S3**).

Among all the aneuploid cells, aneuploidy in Chr X predominated (35 gains and 235 losses, 52.5% of the total events), followed by Chr 21 (48 gains and 21 losses, 13.4% of the total events) (**Supplementary Fig. S6D**). The remaining 60 cells contained 11.6% of the total aneuploidy events (38 gains and 22 losses).

Chromosome 21 had about half of the autosomal aneuploidy events (**Supplementary Fig. S6E**), with more gains than losses. We noticed that Chr 21 aneuploidy occurred unevenly among individuals (**Supplementary Fig. S6F**), as M10 showed a significantly higher ratio of trisomy 21 (2.5% versus 0.2%) than other individuals did. Among all individuals, trisomy 21 occurred more in males than in females (**Supplementary Fig. S6F**), but monosomy 21 occurred equally between males and females (**Supplementary Fig. S6G**).

The aneuploidy occurrence rate for Chr X (270 cells, 1.31% in total) was significantly higher than that of the other chromosomes, contributing to 72.2% of the cells with CNAs >10 Mb (374 cells) (**Supplementary Fig. S6H**). Although all male Chr X aneuploid cells (n = 15) were disomic, the majority of Chr 21 aneuploidy events were monosomic (235 cells, 87.0%) and occurred mostly in females (255 female cells, 15 male cells, **Supplementary Fig. S6H**). Such Chr X loss is prevalent in females aged > 30 years (228 cells), but is rarely discovered in young females (7 cells).

Identifying aneuploidy events in Chr Y was challenging because of its short unique genomic regions (~ 17 Mb). Therefore, we relied on reads counts, instead of copy number estimations, to deduce Chr Y ploidy number, subsequently identifying 115 male cells with Chr Y loss (0.9% of all male cells). To verify the reliability of this assessment, we used reads counts to analyze Chr X and Chr 21 aneuploidy and compared those results with the previous ones (**Fig. 3C–F**). The reads count distribution had a clear linear relationship among monosomic, normal, and trisomic cells, and almost all the results were consistent with the bin-based method (98.37%).

### Loss of heterozygosity and allelic bias in copy number alterations

Using phased, personalized genomic information from F03 and F06 (**Fig. 4A, Supplementary Notes**), we investigated whether somatic CNA events were allele specific. We analyzed all copy neutral chromosomes (CN = 2) in 2,668 cells from F03 and F06 and found only 4 cells (2 from F03 and 2 from F06, 0.1% and 0.2% per individual, respectively) contained copy neutral loss of heterozygosity events in Chromosomes 1, 14, 21, and 22 (**Fig. 4B**). We then focused on CN = 1 genomic regions, within which the allelic pattern should have been either single parental or segmental. Of the 72 CN = 1 events that we analyzed, 70 (97.2%) were single parental and the other two had shuffled genotypes without segmental patterns (**Fig. 4C–E, Supplementary Fig. S7**), likely due to the low probability of collision between a CNA occurrence and recombination.

**Fig. 4.**
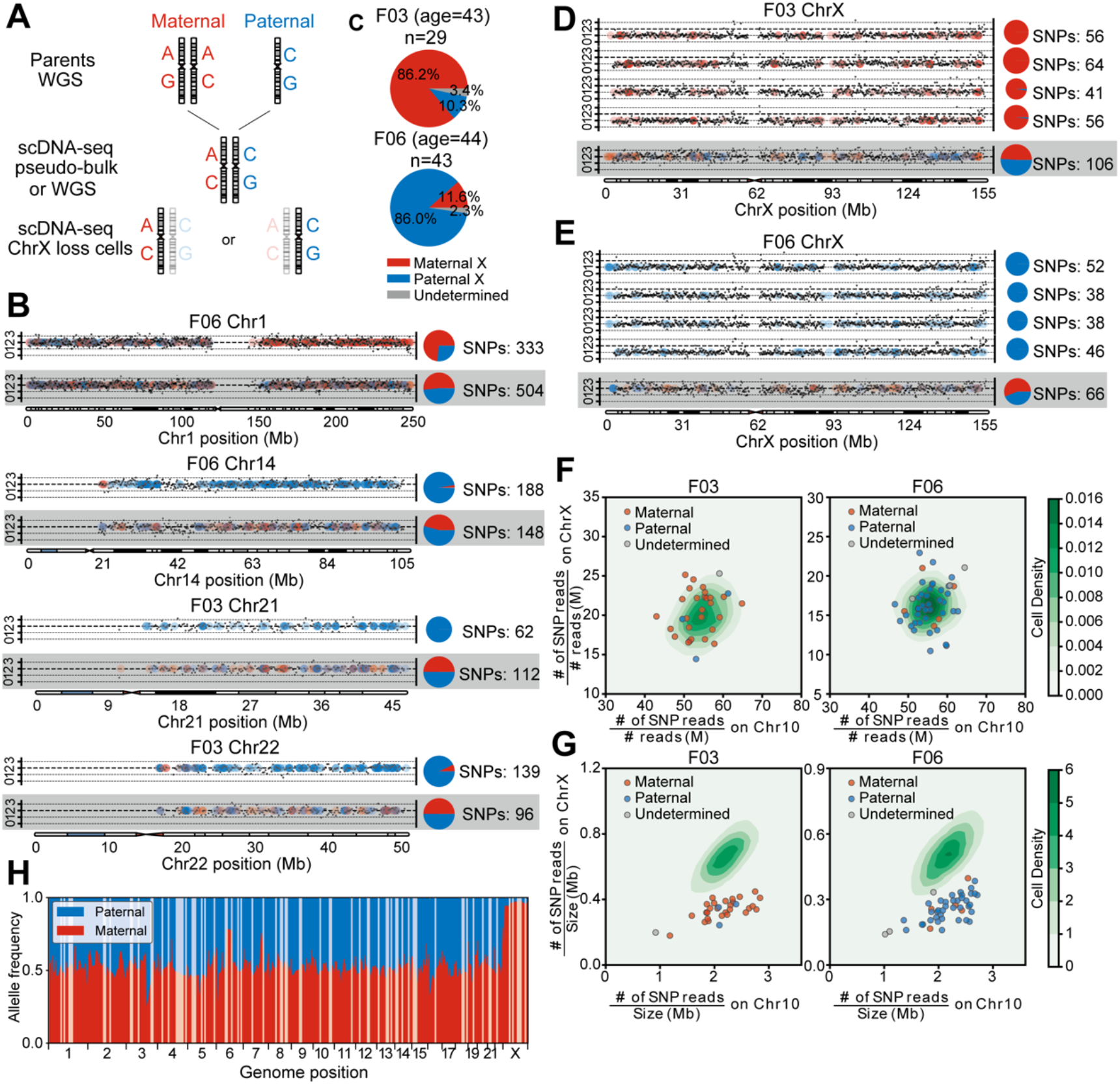
Single-cell (female) haplotype analysis. **(A)** Haplotype identification pipeline of representative chromosome loss cells. For each candidate, two parental genomes were sequenced by whole genome sequencing (WGS). Then, using WGS data or merged single-cell data (pseudo-bulk), we identified each candidate’s heterogeneous sites. Combined with parental data, the heterogeneous sites were labeled paternal or maternal. Finally, we analyzed each candidate’s single cells that had chromosome loss and extracted heterogeneous sites covered by reads, identifying each as paternal or maternal. **(B)** Copy number profiles and haplotype identifications of cells with lost heterozygosity. Black dots show the copy number for each genome locus and dashed lines indicate the integer copy number (0,1,2,3). Colored dots represent the sources of a heterozygous site (red, maternal; blue, paternal) and the pie charts demonstrate the paternal/maternal compositions (numbers of single-nucleotide polymorphisms [SNPs]) of heterozygous sites on each chromosome. Results from normal cells are shown on the bottom for comparison. **(C)** Paternal and maternal allele composition of two individuals’ cells with lost X Chromosomes. Most of the X-loss cells were of the same parental allele. **(D, E)** Copy number profiles and haplotype identifications of cells with lost X Chromosomes and of normal cells, as controls (bottom lines). **(F)** SNP densities and **(G)** reads densities on chromosomes of normal and ChrX-loss cells. The contour plots show the normal cells’ distributions and the scatter plots show the ChrX-loss cells. Cells with lost X Chromosomes that have lower reads density on ChrX, but comparable SNPs density, were not contaminated by male or parental cells. **(H)** Allele compositions of heterogeneous sites in bulk RNA-seq reads of F03 cells. Most of the genomic region displays bi-allelic expression (allele composition ~ 50%), but Chr X clearly shows a maternal bias, which is corrected when most Chr X-loss cells are paternal.

We next examined whether monosomy X cells were parentally biased during chromosome loss, and found most of the F03 monosomic X lymphocytes (86.2%, 25/29 cells) were maternal, while most of the affected F06 monosomic X lymphocytes were paternal (86.0%, 37/43 cells) (**Fig. 4C–E, Supplementary Fig. S7**). To exclude possible technical artifacts and contamination, we calculated the single-nucleotide polymorphism (SNP) density ratio between Chromosome X and Chromosome 10 and found that the ratio distributions for both normal and monosomy X cells were similar (**Fig. 4F**). However, when calculating the normalized reads numbers, we found that the distribution of reads ratio between Chr X and 10 for normal cells was similar, but it was inconsistent for monosomic cells, as only half the reads mapped to Chr X (**Fig. 4G**).

To examine whether parental preference in Chr X loss might correlate with Chr X inactivation, we conducted bulk RNA-seq, based on phased SNPs, to determine F03’s allele-specific expression across her whole genome (**Supplementary Notes**). As expected, autosomal genes expressed unbiased biallelic expression (**Fig. 4H, Supplementary Fig. S7**), but Chr X expression was greatly biased toward the maternal allele, coincident with the fact that the Chr X loss in F03 was mainly paternal (**Fig. 4C, D**).

## Discussion

In this study, we used high-throughput Tasc-WGS to profile CNA landscapes of more than 20,000 single lymphocytes sampled from 16 individuals. Even though CNAs were detected in blood lymphocytes of every donor, the occurrence rate of CNAs >10 Mb for each individual was rather low and correlated little to age. Such low-frequency events can be accurately identified only by profiling thousands of single cells per person; and from the technological perspective, the scalability of single-cell WGS is key to enabling such observations. Additionally, a benefit of large-scale high-performance single-cell sequencing libraries is that we can use them to identify rare CNA events at high resolution, a formerly impractical achievement using other throughput-limited single-cell WGA methods, or by using bulk samples.

We found that CNAs, including the large ones (>10 Mb), were widely distributed throughout the lymphocyte genomes; thus revealing that, on average, about 5% of the lymphocytes of a healthy human have large CNAs. Also, losses were more prevalent than gains, thus suggesting that losses occur more readily than gains do. Previous studies of neurons showed a similarly scattered CNA pattern throughout the genome (Cai et al. 2014; Chronister et al. 2019). Some researchers think that the activity of mobile elements, like that of the active long interspersed nuclear element 1 during brain development, is a major cause of CNAs (Evrony et al. 2015; Sanchez-Luque et al. 2019; Richardson et al. 2014; Erwin et al. 2016). However, unlike that of neurons, the lymphocyte regeneration rate is high, and CNAs harbored in precursor cells may pass to descendent cells via cell differentiation or division. Indeed, most CNA-containing cells are not clonal—we observed only 51 clones in our 20,594-cell sample. In terms of the number of cells, the sizes of these clones were insignificant, thus indicating that rather than being generated in the early developmental stages, they were newly generated. Lymphocytes can expand through mitosis, so some low-frequency CNA events may be inherited.

The CNAs seemed to occur in no particular location across the genome, except in Chromosomes 21, X, and Y, where most of the aneuploidies were found. Most segmental CNAs were randomly scattered across the whole genome, but whole-chromosome CNAs likely affected many genes, either by completely silencing or altering their expression levels (Zhang et al. 2009; Girirajan et al. 2011). As a result, aneuploidies are lethal in most cases (Hassold et al. 2001; Santaguida et al. 2015). However, 2.4% of the cells in our study had aneuploid events, especially at Chr 21, which displayed trisomy, the most prevalent human aneuploidy (Sanchez-Luque et al. 2019; Richardson et al. 2014). This consistency suggests similar selection outcomes for aneuploid events in both humans and lymphocytes.

Aneuploidy occurs more in sex chromosomes than in autosomes. Although our results showed a weak correlation of age with CNA occurrence, monosomy X is rare in young females but becomes more prevalent in females over age 30, for whom the rate can reach 2%–7%. Such a chromosome loss was parentally allele specific in the affected individuals. According to haplotype information and bulk RNA-seq data, we found that most lost X Chromosomes were inactive (Xi). These results agree with previous studies that suggested that the X inactivation skewness pattern is more prevalent in older than in younger individuals (Sharp et al. 2000; Sandovici et al. 2004; Amos-Landgraf et al. 2006; Machiela et al. 2016; Zito et al. 2019). This association may be due to the lethality of a lost active X, so most monosomy X cells are missing Xi because negative selection assumes a randomly generated loss during mitosis. Unlike the high incidence of monosomy X cells, trisomy X cells are rarely found. This imbalance further suggests that Chromosome X aneuploidy likely does not result from simple uneven separation during mitosis. Additionally, compared with autosomal aneuploidies, Xi loss is both non-lethal to affected cells and under positive selection. Therefore, we observed a higher incidence of monosomy X cells than cells with autosomal events. Another possible contributing factor is that conformational changes to Xi may ease its loss during mitosis (Galupa et al. 2018). Obviously, more investigation is needed to clarify the mechanisms of these phenomena.

Our results demonstrate how shallow WGS, after extending throughput to 1,000 single cells, enables quantitative identification of rare copy number change events. However, the sensitivity of CNA detection was limited in this study because, although it performs better than most other existing methods, Tasc-WGS harbors intrinsic coverage noise due to amplification bias and coverage stochasticity. To lower that false positive CNA identification rate, we applied strict criteria to screen such events, even though it caused us to lose some sensitivity and resolution. Studies of cancers (Beroukhim et al. 2010; Navin et al. 2011; Jacobs et al. 2012; Laurie et al. 2012) and neuronal disorders (van den Bos et al. 2016; Yurov et al. 2007; Bundo et al. 2014) would benefit from the ability to identify smaller CNAs, but improvements of both experimental protocols and computational algorithms are needed. For instance, the commonly used data processing pipelines for determining copy number are based on bulk sequencing or micro-array data, not single-cell data. With the popularity of single-cell profiling and the availability of more data, such as ours, a need for more appropriate computational approaches must be met soon.

## Methods

### Ethics approval

This study was approved by the Ethics Committee of Tsinghua University (No. 20180011), Ethics Committee of the Cancer Hospital, the Chinese Academy of Medical Sciences and Peking Union Medical College (No. NCC2017G-002), and the Ethics Committee of Fuwai Hospital, Chinese Academy of Medical Sciences and Peking Union Medical College (No. 2017-880).

### Patients and clinical samples

We recruited Fuwai Hospital patients who had cardiovascular diseases, as well as their visiting family members. A single patient with colon cancer was enrolled through the Cancer Hospital. Patients and family members were given full research program descriptions, which included potential risks. We obtained informed consent from all patients and family members before genetic testing, and then collected fresh blood samples from both healthy donors (M01-M03, F03-F06) and cardiovascular disease patients (M04-M10, F01, F02), as well as a tumor sample from the cancer patient after surgery.

### Peripheral blood mononuclear cell isolation and single-cell sorting

We used Ficoll-Paque PLUS (Cytiva, #17-1440-02) according to the manufacturer’s instructions to isolate mononuclear cells from fresh blood samples. Briefly, for each sample, Ficoll-Paque medium (3 ml) was added to a 15-mL centrifuge tube and then a blood sample (2 mL) diluted 1:1 in phosphate buffered saline (PBS) was carefully layered onto the Ficoll-Paque medium. The tube was then centrifuged at 400*g* for 30 min at room temperature. The second layer, which contained mononuclear cells, was pipetted out, transferred to a new tube, and washed twice in 10 mL PBS before being resuspended in 1 mL PBS-bovine serum albumin (BSA) buffer. Typically, we isolated 1 × 10^6^ cells from each sample. We then used a FACSAria III sorter (BD Biosciences), gated for lymphocytes and singlets, to sort out single cells according to forward and side scatter signals. We then placed each sorted, single cell directly into 2 µl lysis buffer (30 mM Tris-HCl [pH = 8.0], 10 mM NaCl, 0.2 µL Proteinase K [Qiagen, #19133], 5 mM EDTA, and 0.5% Triton X-100 [Sigma-Aldrich, #T9284]) in a well of a 96-well plate.

### Single-cell isolation from a tumor tissue

We ground the colorectal cancer sample (~ 0.1 cm^3^) using a dounce glass tissue grinder. The cells were then washed, resuspended in PBS, and filtered through a Falcon 40-µm cell strainer. They then underwent fluorescence-activated cell sorting (FACS), gated for single-cells, and each cell was sorted into a well in a 96-well plate.

### Culturing and isolating single cells and optimizing cell lines

We used GM12878 cells (Coriell Institute) and HEK293 cells (American Type Culture Collection) for protocol optimization. Those cells were cultured at 37 °C under 5% CO_2_ in a humidified incubator. We cultured GM12878 cells in RPMI 1640 medium (Gibco, #C11875500BT) with 10% fetal bovine serum (Gibco, #10100147) and 1% penicillin–streptomycin (Gibco, #15140122), then spun the cell suspension at 500*g* for 5 min, discarded the supernatant, and washed the cell pellet twice using PBS before resuspending it in PBS with 1% BSA. We cultured HEK293 cells in DMEM medium (Gibco, #11965092) with 10% fetal bovine serum and 1% penicillin– streptomycin. The cells were then washed twice using PBS, detached by adding 1 mL 0.25% trypsin-EDTA (Gibco, #25200056) to their culture dish, centrifuged at 500*g* for 5 min, and recovered in 1% PBS-BSA buffer. All cells underwent FACS that was gated for single-cells and each cell was subsequently sorted to a well in a 96-well plate.

### Purification of genomic DNA

We purified genomic DNA (gDNA) using a Genomic DNA Purification Kit (Thermo Fisher Scientific, #K0512) according to the manufacturer’s instructions. We then quantified that DNA with a Qubit fluorometer system (Invitrogen) and diluted it to 6 pg/μL.

### Single-cell whole genome amplification and sequencing

The 96-well plates were then centrifuged at 2,000*g* for 1 min and a lysis reaction proceeded at 50 °C for 3 h. We added tagmentation buffer (1x TD buffer, 0.015 µL TTE Mix V50 [Vazyme, #TD501], 0.625x protease inhibitor cocktail [Promega, #G6521], and 1 mM MgCl_2_) to reach a volume of 10 µL per well and then incubated the plates at 55 °C for 1 h. Tagmented DNA fragments were amplified by adding 12 µL PCR master mix composed of 11 µL Q5 High-Fidelity 2x Master Mix (New England Biolabs, #M0492) and 0.5 µL each of 10 mM Nextera i5 and i7 index primers. PCR thermocycling conditions were 72 °C for 8 min, 98 °C for 30 s, 24 cycles of 98 °C for 15 s each, 60 °C for 30 s, and 72 °C for 90 s, with a final incubation at 72 °C for 5 min. The subsequent PCR products were merged in groups of 5 plates (480 single-cell wells) and then purified using 1x VAHTS DNA clean beads (Vazyme, #N411). Library quality control was conducted on a 5200 Fragment Analyzer System (Agilent, #M5310AA) to determine fragment distribution, and then qualified libraries were quantified and sequenced on a HiSeq X Ten System (Illumina) following the manufacturer’s standard protocols.

### Bioinformatic analyses

#### Data processing

Paired-end reads were aligned to the human reference genome (hg38) using nvBowtie (https://github.com/NVlabs/nvbio), a graphics processing unit-accelerated version of Bowtie 2 (Langmead et al. 2009). Then, each cell’s mapped reads were demultiplexed using perfectly matched cell barcodes. Typically, 0.3 million reads were sufficient for copy number profiling at a 200-kb resolution. Before downstream analysis, we excluded cells with less than 0.3 million reads, keeping reads mapped with minimum mapping quality scores of 20, and removed PCR duplicates using SAMtools (Li et al. 2009).

#### Copy number profiling and quality control

We applied two methods, HMMcopy (Shah et al. 2006) and DNAcopy (Olshen et al. 2004), to calculate the copy number profiles of each sample at the 200-kb resolution, with GC content and mappability normalized. Both algorithms are commonly used in single cell studies, but they each give different identification results for small size variations (Knouse et al. 2016). HMMcopy uses a hidden Markov model (HMM) to determine copy number, while DNAcopy applies circular binary segmentation (CBS) for analysis. We combined the two methods to further increase the specificity and accuracy of CNA identification.

We used the Bayesian information criterion as a metric to evaluate model fitness with different computational parameters in HMMcopy and DNAcopy calculations, using the strictest parameters (alpha = 10^−4^ for DNAcopy and e = 0.9999 for HMMcopy) under the same fitness to enhance CNA calling specificity.

After segmentation, we used three features to assess the quality of the single cell sequencing results, and then filtered out low-quality cells and incorrect segmentation calls. First, we checked the average of all copy numbers identified in each bin (degree of ploidy abnormality), and that value was greatly influenced by cell ploidy. Cells with abnormal ploidy at the whole genome level (ploidy > 3) were discarded. We then checked the median absolute pairwise difference (MAPD), typically used for indicating amplification evenness, to rule out poorly amplified cells (MAPD > 0.6). Finally, we checked each cell’s number of segments. We noticed that some cells exhibited acceptable MAPD values but had fragmented copy number profiles. This could have been caused by incomplete lysis, contamination from other cells or cell debris, or during the S-phase, as some studies have suggested (Laks et al. 2019; Chen et al. 2017). Since cells with a CNA or a fragmented chromosome will have more segments and slightly higher MAPD values than would normal cells, the other genomic regions of those cells are still high quality. So, we then calculated the number of segments (degree of fragmentation) and MAPD for each chromosome and used the third highest values to represent each cell’s value.

#### Identification of copy number alteration events

We identified each CNA by combining the two algorithms, CBS and HMM, and keeping the double-positive counts as true events. Since both algorithms are sensitive to the local contents of copy number profiles (Knouse et al. 2016; Zhang et al. 2015), especially for losses, we developed a shuffling pipeline to improve the confidence of identifying CNA events.

For each cell with CNAs, segments with amplification or deletion were shuffled throughout the genome and were re-identified by CBS and HMM algorithms. We identified CNA events with high confidence by repeating the shuffle process 20 times and averaging the copy number values identified for a given shuffled segment. A loss was defined as a segment with a copy number value less than 1.4. Only those CNAs larger than 2 Mb were kept for downstream analysis.

To avoid false identification affected by mapping uniqueness, we ruled out those CNAs either located near centromeres (overlapping more than 40%) or with disperse copy number profiles (with larger deviations [mean or median > 0.4]) between the copy number values of bin and segment, usually at the chromosome ends).

#### Simulation of CNA profiles

Since bins of CNA profiles are normally distributed (Nilsen et al. 2012), we generated normally distributed simulated CNA data to investigate false positive (FP) calls introduced by the algorithm. We set σ values to range from 0.4 to 0.8 and then generated 1,000 simulated CNA profiles for each σ, replicating the process three times. We then adopted the same CNA calling pipelines and counted the FP CNA events in each batch.

#### Estimating the coefficient of variation of CNA identification

Large CNAs are rare events and vulnerable to sampling errors. Therefore, we simulated the sampling process by sampling different numbers of cells (sample size, from 3 to 10^5^) from 10^6^ cells having different ratios of CNA-containing cells (from 0% to 20%). We repeated each test 100 times to determine sufficient sample size and then calculated the coefficient of variation (CV) for each condition.

#### Clone identification

We first used dimension reduction to view all the cells with >10 Mb of CNAs. First, CNA profiles having 200-kb resolutions were smoothed using a 4-Mb window; and then, using multidimensional scaling, they were transformed into a low-dimensional representation. We adjusted the number of dimensions representing each chromosome (each dimension represents ~ 10-Mb CNA profiles) and concatenated all chromosomes. Then, we were able to visualize low-dimensional representations of CNA profiles in 2-dimensions by using *t*-stochastic neighbor embedding.

We identified clonal CNAs by calculating the relative CNA ratio in every chromosome for every individual. Specifically, for each individual, we calculated either the fraction or the number of CNAs in each chromosome to represent the enrichment of CNAs in each, and then normalized those values based on chromosome lengths.

To further investigate the clonal CNAs, we calculated the Euclidean distance between cells for every individual and identified similar cell pairs according to their distance distributions. We then constructed an undirected graph using cells as nodes and the Euclidean distances as edges and identified clones as maximally connected subgraphs.

#### Haplotype calling

We first genotyped genome sequencing data on all loci of the whole genome with the same pipeline. Reads were first trimmed and filtered using the following criteria. The adaptors were removed according to the reverse complementary sequence of the paired-end reads, and filtered reads were dynamically trimmed with a Phred cutoff of 20. The remaining reads were then mapped to the human reference genome using Bowtie 2 (MapQ ≥ 40, XM < 4), and whole genome genotypes were called using the UnifiedGenotyper mode of GATK-3.5 (DePristo et al. 2011). We performed heterozygosity analysis with a minor allele frequency cutoff between 30%–50% and with 0%–20% homozygosity. Variant call format (VCF) files of three sample genotypes were merged into one VCF file, and heterozygous loci of those three samples were extracted into a locus file as a union for VCF scanning. Only those loci from which either the mother is heterozygous and father homozygous or the father is heterozygous and mother homozygous were used to phase the child’s haplotype.

#### Analysis of Chr X

For each single cell from F03 and F06, reads with single-nucleotide polymorphisms (SNPs) were identified using SAMtools (base quality > 30). Then, the haplotype for each SNP was labeled as paternal, maternal, or neither (likely due to sequencing error) using the haplotype map. Haplotype counts for each bin were the sums of every SNP site in that bin. For monosomy X cells, SNPs in every bin within segments that identified a loss, were summarized and identified as paternal, maternal, or undetermined (binomial test, *p* < 0.001).

#### Analysis of Chromosome Y

Since Chr Y had few uniquely mapped reads, we had to develop a special method to determine its copy number. Reads coverage of Chr 21, X, and Y were calculated by SAMtools normalized by sequencing depth, and then cells with Chr 21 and Chr X aneuploidies were identified by coverage depth. If we calculated the percentage of sequenced reads belonging to Chr 21 or Chr X of each single cell, we could also easily identify cells with normal or altered copy numbers in those two chromosomes. Actually, we found that more than 98% of those results were consistent with the results determined by coverage depth. We then applied our percentage classification method to identify the Chr Y copy number for each single cell.

### Data access

The whole-genome and RNA-seq data generated in this study are deposited in the Genome Sequence Archive (GSA; https://ngdc.cncb.ac.cn/gsa-human) in National Genomics Data Center, China National Center for Bioinformation / Beijing Institute of Genomics, Chinese Academy of Sciences under accession number HRA001513. The scripts generated for the bioinformatics analysis are available in the Supplemental Code.

## Acknowledgements

We thank Chenyang Geng from the Peking University High-throughput Sequencing Center and Biomedical Pioneering Innovation Center, and Fei Wang from the National Center for Protein Sciences (Peking University) for the experimental assistance. This project was supported by National Key R&D Program of China (2018YFA0108100 to Y.H.), National Natural Science Foundation of China (22050002 to Y.H.), Beijing Municipal Science and Technology Commission (Z201100005320016 to Y.H.), Beijing Advanced Innovation Center for Genomics, and Shenzhen Bay Laboratory.

## Author Contributions

Y.H., J.W., and Z.Z. conceived the study; L.L., H.C., J.Z., L.D., J.S., and S.G. performed experiments; H.C., L.L. and M.D. performed data analyses; Y.F., L.L, L.D., and H.C. developed the Tasc-WGS protocol; C.S., J.W., and Y.L provided clinical samples; L.L., H.C., Y.H., and J.W. wrote the manuscript with inputs from all authors; Y.H., J.W., and Z.Z. supervised all aspects of this study.

