## Supplementary_Material for "Low frequency somatic copy number alterations in normal human lymphocytes revealed by large scale single-cell whole genome profiling"

[1] Biomedical Pioneering Innovation Center (BIOPIC) and Beijing Advanced Innovation Center for Genomics (ICG), School of Life Sciences, Peking University, Beijing 100871, China.
[2] National Clinical Research Center of Cardiovascular Diseases, State Key Laboratory of Cardiovascular Disease, Fuwai Hospital, National Center for Cardiovascular Diseases, Chinese Academy of Medical Sciences and Peking Union Medical College, Beijing 102300, China.
[3] Department of Oral Pathology, Peking University School and Hospital of Stomatology, National Engineering Laboratory for Digital and Material Technology of Stomatology, and Beijing Key Laboratory of Digital Stomatology, Beijing 100081, China.
[4] School of Life Sciences and Beijing Advanced Innovation Center for Structural Biology (ICSB), Tsinghua University, Beijing 100084, China.
[5] Department of Neurobiology, Capital Medical University, Beijing100069, China.
[6] Department of Etiology and Carcinogenesis, National Cancer Center/Cancer Hospital, Chinese Academy of Medical Sciences and Peking Union Medical College, Beijing 100021, China.
[7] College of Chemistry and Molecular Engineering, Beijing National Laboratory for Molecular Sciences, Peking University, Beijing 100871, China.
[8] Peking-Tsinghua Center for Life Sciences, Peking University, Beijing 100871, China.
[9] Institute for Cell Analysis, Shenzhen Bay Laboratory, Shenzhen 518132, China.
[#] These authors contributed equally to this work.

### Supplementary Notes

Validation experiments

*Using diluted genomic DNA to construct sequencing libraries*

To evaluate the experimental protocol's reproducibility and technical specifications, we constructed a number of sequencing libraries from mock single cells using diluted genomic DNA. Genomic DNA extracted from the M01 blood sample was diluted, mixed with lysis mix, and then aliquoted into a 96-well plate containing tagmentation mix. In each well, 6 pg gDNA was aliquoted, indexed, and amplified by PCR, then all 96 samples were merged and sequenced on a HiSeq 4000 system (Illumina).

*Examining cross-contamination by mixing human cells with mouse cells*

To evaluate possible contamination between different wells during high-throughput single-cell library preparation, we first constructed an experimental library containing both cells from a mouse cell line, 3T3, and HEK293 cells. We sorted single 3T3 cells into the odd-numbered column wells of two 96-well plates, and single HEK 293 cells into the even-numbered column wells, for a total of 192 cells. We then proceeded with the same single-cell whole genome amplification and sequencing process described above. After demultiplexing by cell index, reads were aligned to the human reference genome (hg38) and mouse reference genome (mm10). We then calculated each sample's mapping rates on both reference genomes to identify the cell type in each well.

*Fluorescence in-situ hybridization*

We conducted fluorescence in situ hybridization, at the Beijing Hightrust Diagnostic Laboratory, to identify copy number change events in the M01, M02 and F03 blood samples. We used probes CEP3 (D3Z1), CEP8 (D8Z1), RB1, CEP18 (D18Z1),

D21S259/D21S341/D21S342, and CEPX (DXZ1) to characterize copy number alternations (CNAs) in Chromosomes 3, 8, 13, 18, 21, and X, respectively. Detailed statistics can be found in **Table S3**.

### Supplementary Figures

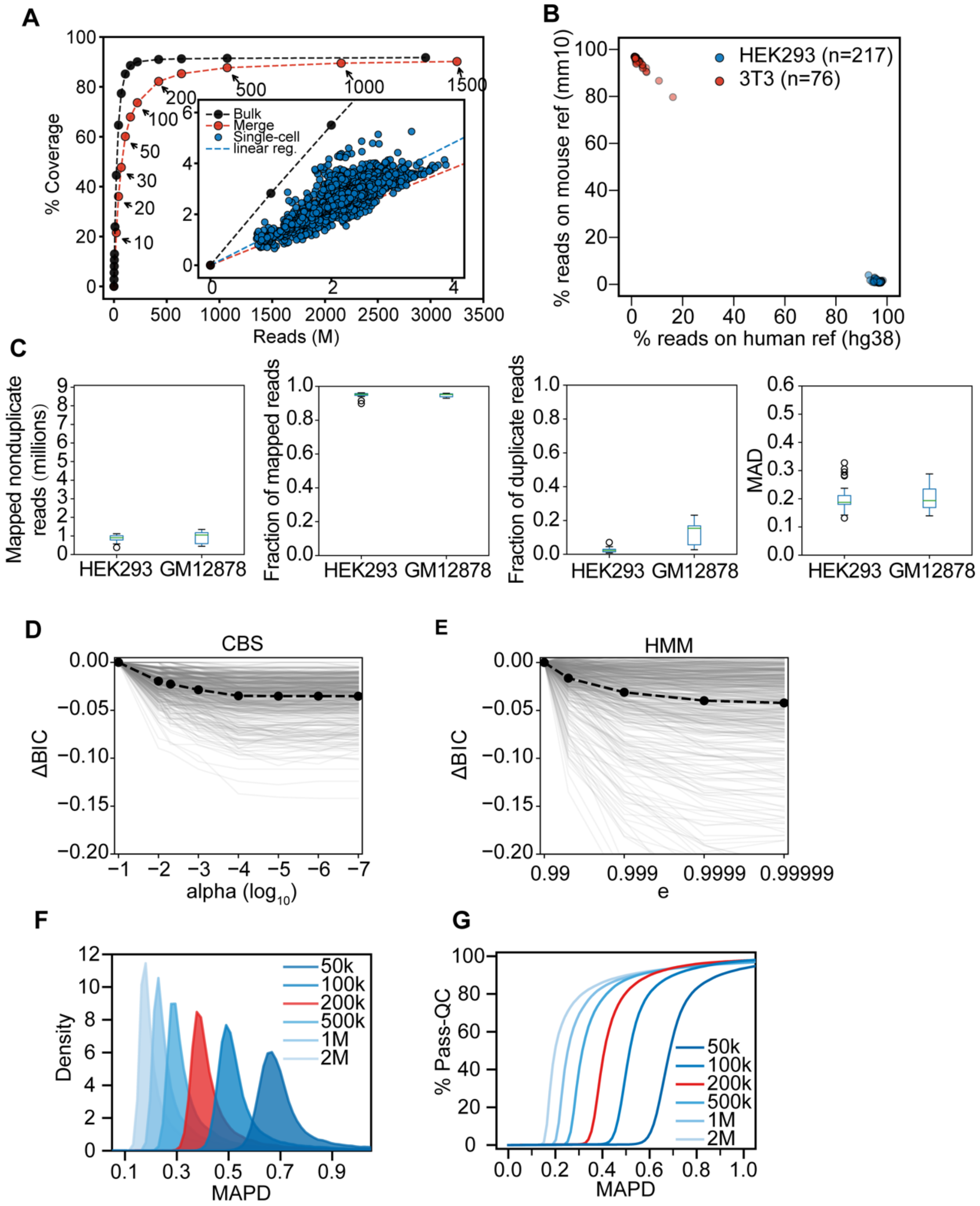


**Supplementary Fig. S1. Quality evaluation of the single-cell data. (A)** Genome coverage analysis. All the single-cell data (blue dots) show unified reads numbers and coverage. The blue dashed line indicates the linear correlation between reads and coverage. Compared to bulk data (black dots and dashed line), single-cell coverage is slightly lower under comparable reads, but once enough cells aggregate (red dots and dashed line), coverage is comparable to that of the bulk data. **(B)** The cross-species experiment showed that cross contamination between wells in the plates was negligible, as mouse cells mapped to the mouse genome and human cells mapped to the human genome. **(C)** Sequencing metrics of HEK293 and GM12878 single-cell samples. MAD: median absolute deviation of bins assigned to a 2-copy state. Boxes show the 25$^{th}$ and 75$^{th}$ percentiles and the horizontal bars show means, whiskers show min/max values, and circles are outliers. **(D, E)** Parameter optimization of the **(D)** circular binary segmentation (CBS) and **(E)** hidden Markov model (HMM) algorithms. We used Bayesian information criteria (BIC) to evaluate which parameter fit the CBS and HMM algorithms best. Grey lines indicate results in different cells and black dots with the dashed lines connect the means of 500-cell results. alpha is the significance levels (used in DNAcopy) and e is the probability of extending a segment (used in HMMcopy). **(F, G)** The median absolute pairwise difference (MAPD) distribution under **(F)** different bin sizes and **(G)** different percentages of passing cells (Pass-QC) when using different bin sizes and MAPD cut-offs. In this study, we used a 200-kb bin and 0.6 as the MAPD cut-off for a balanced trade-off (red distribution and line) between resolution and quality.

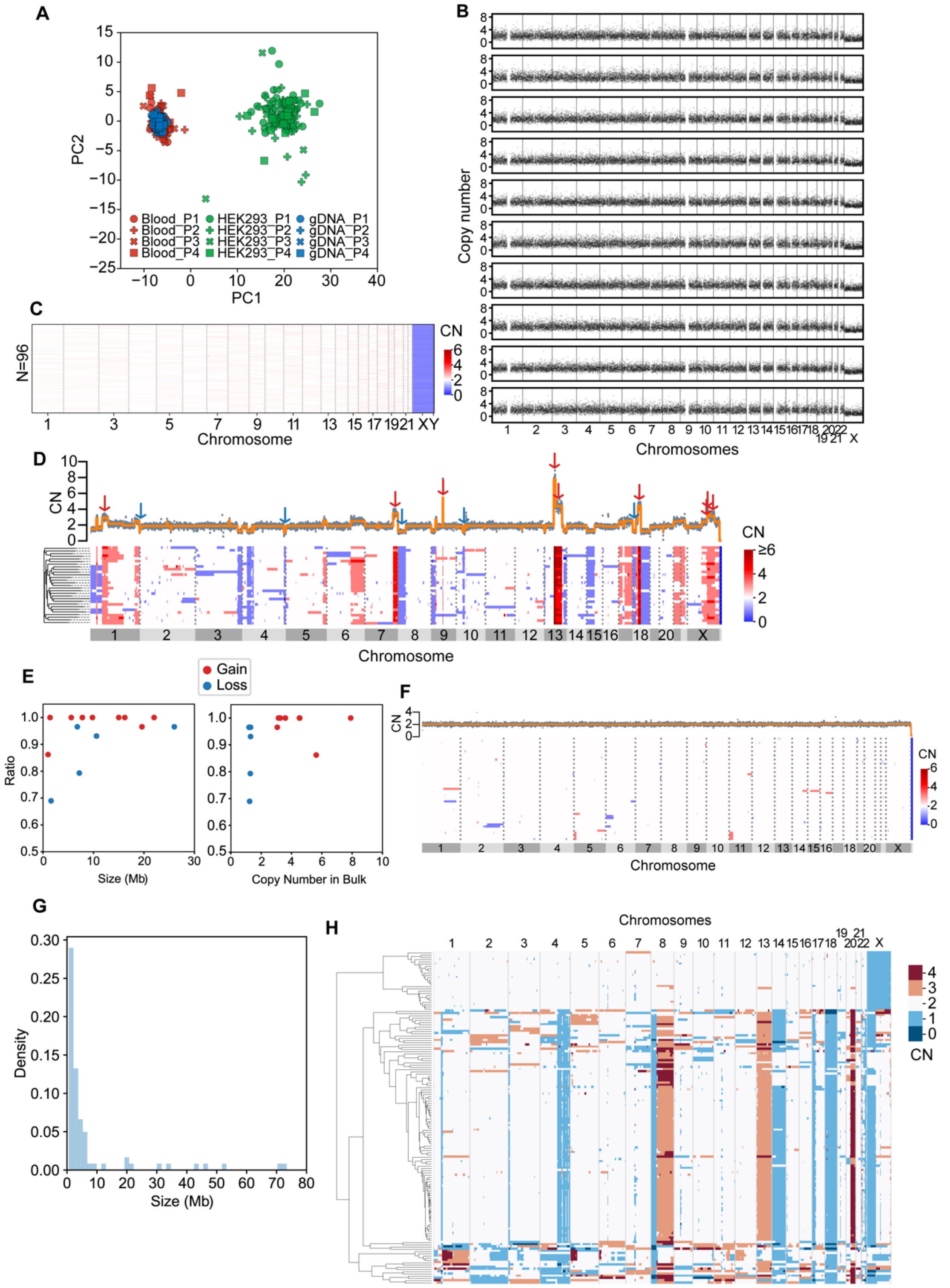


**Supplementary Fig. S2. Tn5-assisted single-cell whole genome sequencing in different samples**. **(A)** PCA of blood cell (red), genomic DNA (gDNA, blue) and HEK 293T cell (green) data, using each 96-plate technical replicate (n = 4) that is represented by its own shape in the graph. **(B)** Ten representative copy number profiles of bulk diluted 6-pg gDNA samples. No large CNAs and aneuploidies were found. **(C)** Summary of the copy number (CN) profiles of 96 bulk diluted 6-pg gDNA samples. **(D)** Copy number profiles of a HEK293 dataset including both one bulk (upper panel) and 29 single-cell samples (lower panel). Fourteen regions in the single-cell dataset were used to calculate detection rates, including copy number gains (red arrows, CN ≥ 3, range 3.06–7.91, median = 3.60) and copy number losses (blue arrows, CN ≤ 1.3, range 1.20–1.30, median = 1.28). **(E)** Scatter plots showing the relationships between the CNA detection ratios and the CNA sizes (left panel) and the copy numbers (right panel) in the HEK293 cell line bulk dataset. **(F)** Copy number profiles for the GM12878 dataset, including both bulk sample (upper panel) and 49 single-cell samples (lower panel). **(G)** The size distributions of CNAs detected in the 49 GM12878 single cells. **(H)** The copy number alteration pattern and clustering of the colorectal cancer sample.

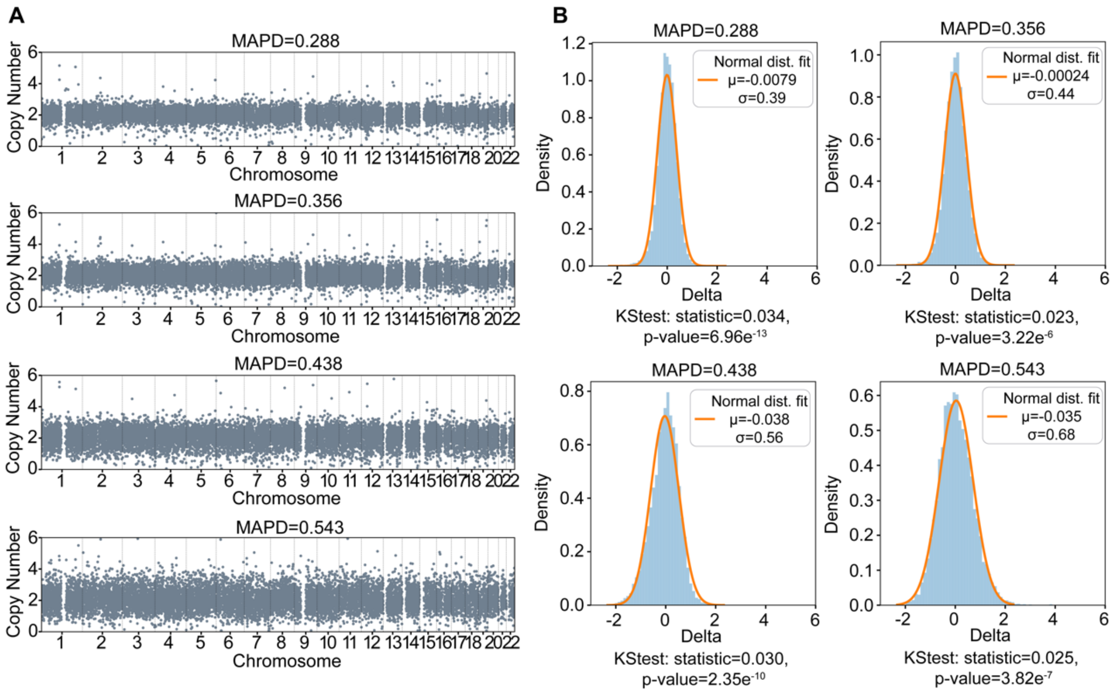


**Supplementary Fig. S3. (A)** Representative examples of CNA profiles of four lymphocyte cells and **(B)** the deviations of their bin values from a neutral value (CN = 2) fitted to normal distributions. MAPD, median absolute pairwise difference; KStest, Kolmogorov-Smirnoff goodness-of-fit test.

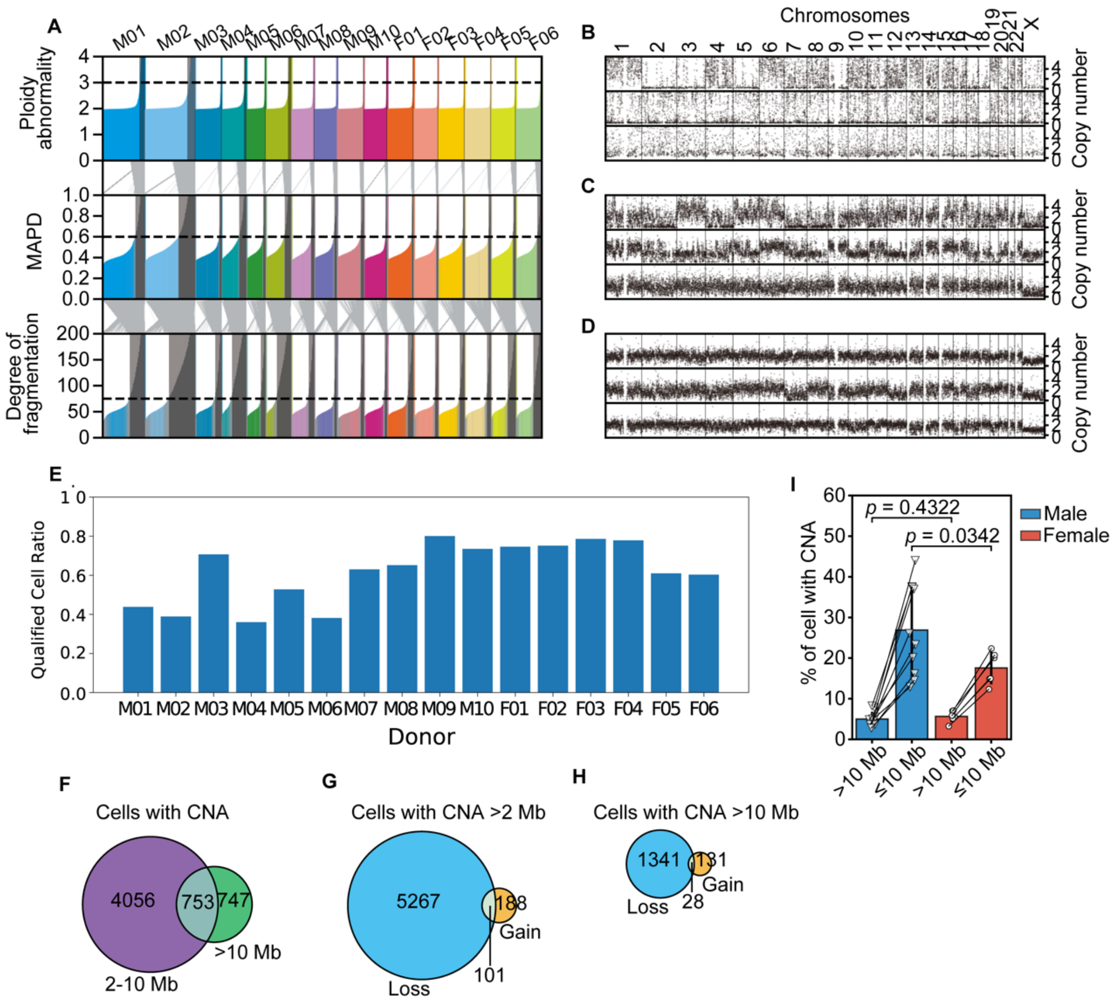


**Supplementary Fig. S4. Data quality filters and basic analyses. (A)** Quality filters for all samples in our study. The histograms show the distributions of three quality criteria. Gray areas identify cells that failed the corresponding filter and gray lines between the histograms follow those cells between criteria. **(B–D)** Copy number profiles of representative cells that failed the three filters in (**A**). The top profiles show cells with abnormal ploidy, middle profiles show cells with large median absolute pairwise differences (MAPD), and bottom profiles show cells with fragmented segments. (**E**) Qualified cell ratios of samples from 16 donors. **(F–H)** Venn diagrams of the numbers of **(F)** cells with 2–1-Mb and/or >10-Mb copy number alterations (CNAs), **(G)** gain and/or loss cells with >2-Mb CNAs, and **(H)** gain and/or loss cells with >10-Mb CNAs. **(I)** Autosomal CNA percentages between male and female samples showing no significant difference between male and female CNA (>10-Mb) percentages (*t*-test).

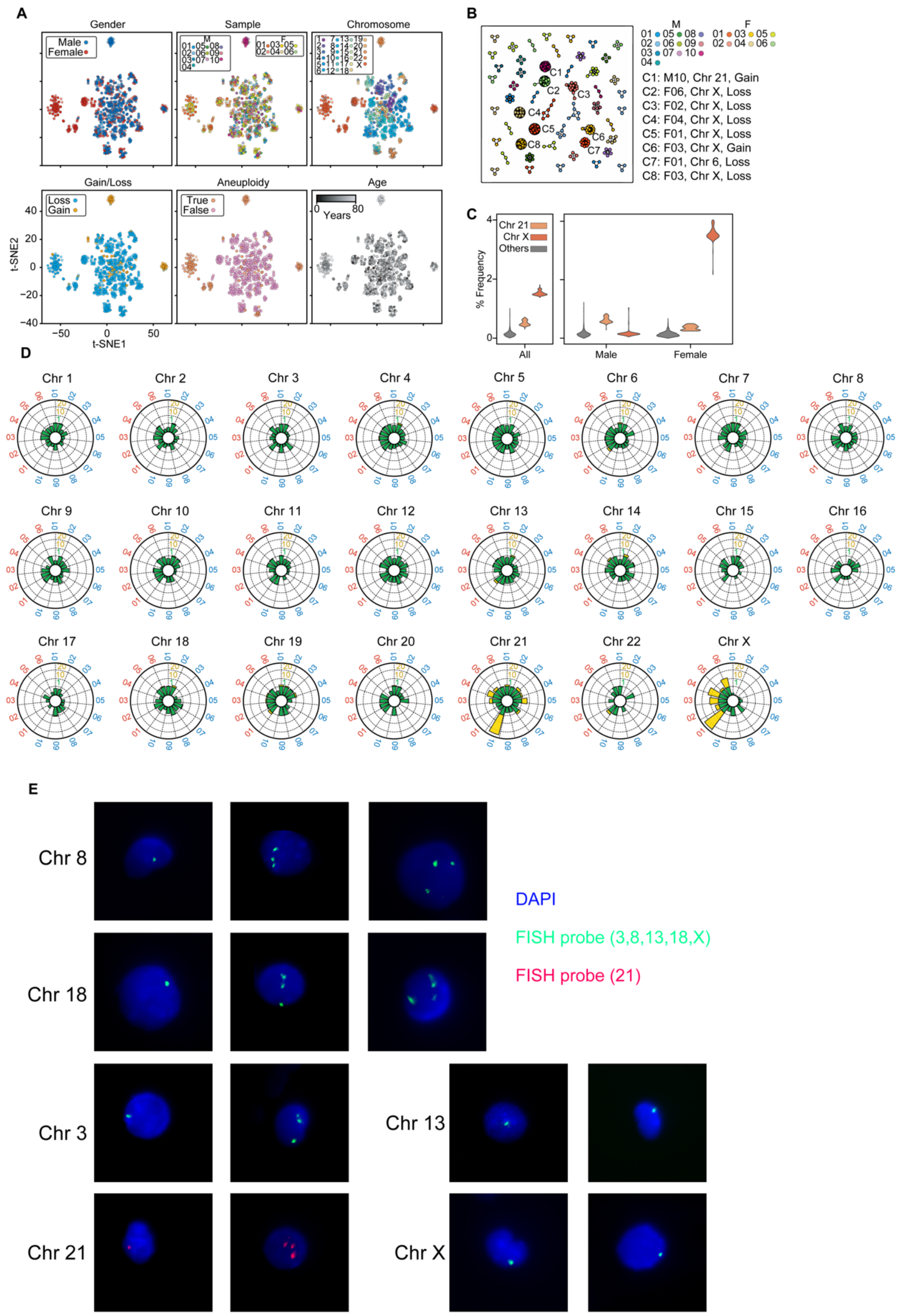


**Supplementary Fig. S5. Clone analysis. (A)** Graphical visualizations of 2-dimensional representations produced through multidimensional scaling of the copy number profiles of cells with >10-Mb copy number alterations (CNAs). t-SNE, *t*-distributed stochastic neighbor embedding. **(B)** All 51 clones shown in an undirected graph with node colors representing cells from different samples and edges representing cells paired with small Euclidean distances, <10. **(C)** CNA frequency in genomic loci (200-kb resolution) of all samples (left) and in male and female samples (right). **(D)** Radar plots demonstrating normalized CNA counts for all chromosomes in each individual, labeled outside each circle in blue for males and red for females. Colors within each plot indicate the counts scale (green for 0–1, yellow for 1–20). **(E)** Fluorescence in situ hybridization (FISH) photomicrographs of cells with aneuploidies. Blue (DAPI) represents the nuclei, green shows the FISH probe on Chromosomes 3, 8, 13, 18, and X, and red shows the FISH probe on Chromosome 21.

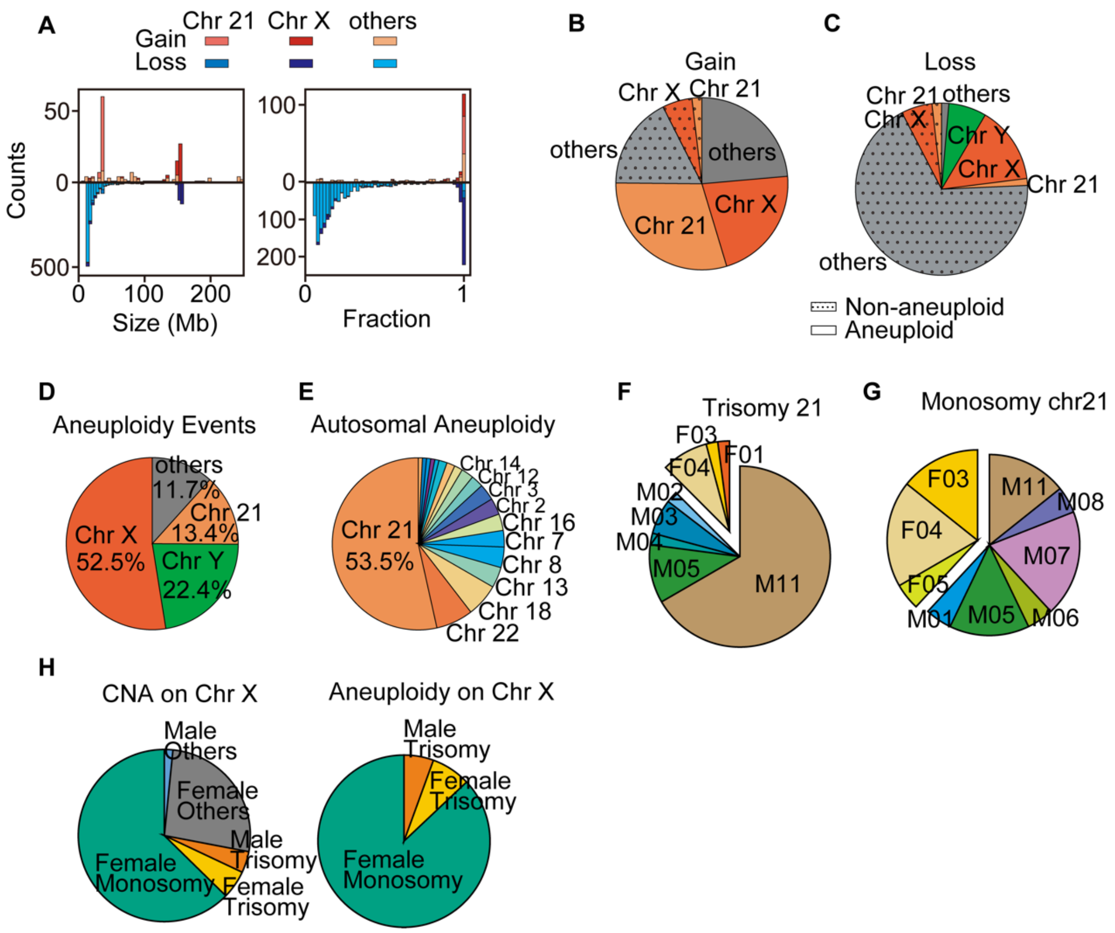


**Supplementary Fig. S6. Aneuploid events and chromosomes. (A)** Size distributions and fractions of >10-Mb gain and loss copy number alterations (CNAs) (excluding Chromosome Y). **(B, C)** Compositions of aneuploid and non-aneuploid CNAs of >10 Mb each: **(B)** gains and **(C)** losses. **(D, E)** Chromosomal compositions of **(D)** all aneuploidy and of **(E)** autosomal aneuploidy. **(F)** Trisomy 21 in male (M) and female (F) samples. **(G)** Monosomy 21 in male and female samples. **(H)** The compositions of all CNAs and aneuploidies on Chromosome X.

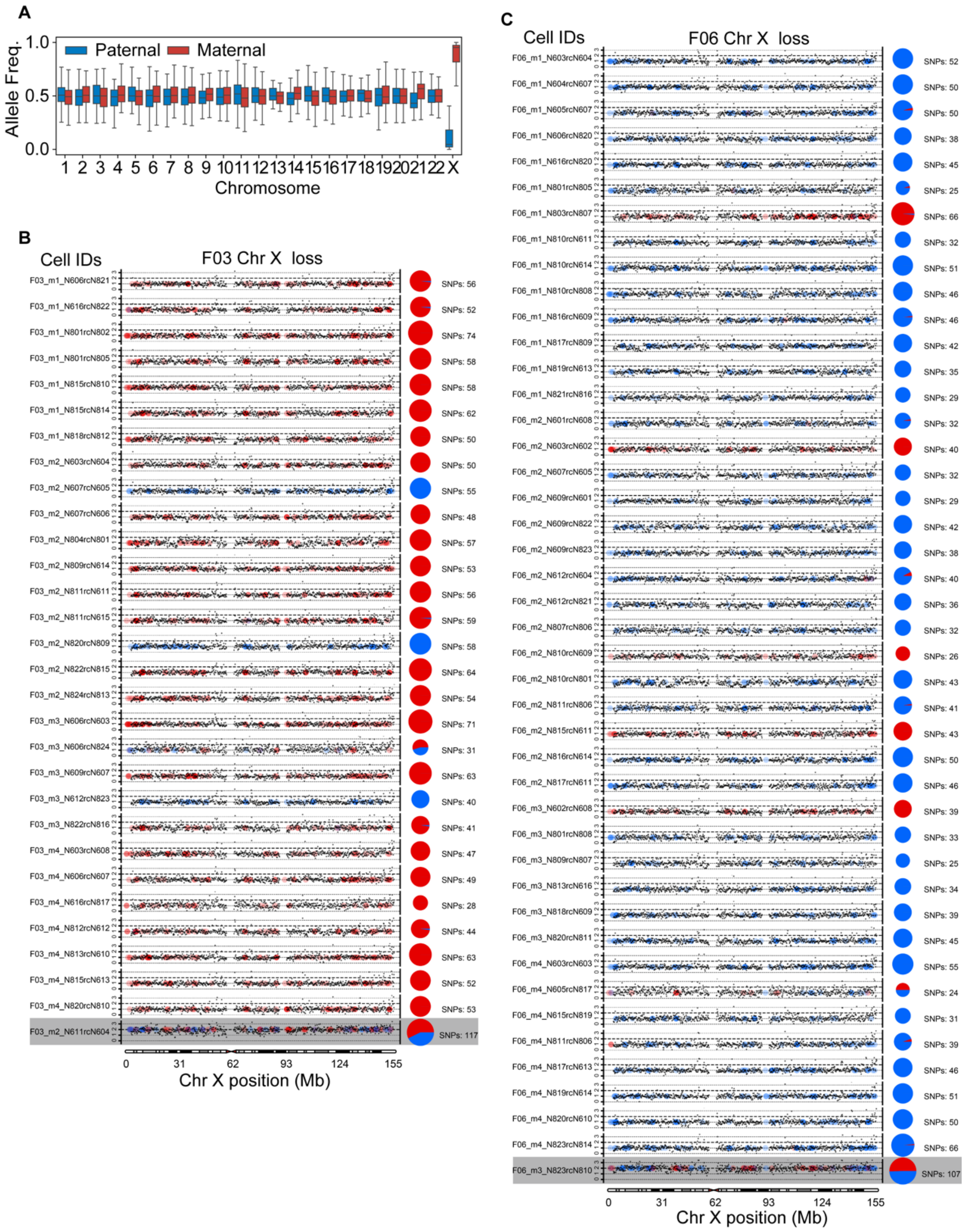


**Supplementary Fig. S7. Chromosome X haplotype analysis**. **(A)** RNA-seq reads of allele frequencies on all chromosomes. **(B, C)** Copy number profiles and haplotypes of all cells from **(B)** F03 and **(C)** F06 with Chromosome X loss. Black dots show the copy number for each genome locus and dashed lines indicate the integer copy number (0,1,2,3). Colored dots represent the sources of heterozygous sites (red, maternal; blue, paternal) and the pie charts show paternal/maternal compositions (numbers of single-nucleotide polymorphisms [SNPs]) of heterozygous sites on each chromosome. Results from normal cells are shown on the bottom for comparison.

**Table S1. Summary of donor individuals and their sequencing results**

| Donor ID | Age | Sex | Disease[1] | Read number (M) | | Cell number | | | /w CNA (% cell) |
|---|---|---|---|---|---|---|---|---|---|
| | | | | Total | Per cell[2] | Input | Pass QF[3] | /w CNA[4] | |
| M01 | 44 | M | — | 6063.26 | 2.14 | 3360 | 1681 | 656 | 39.02 |
| M02 | 33 | M | — | 6832.00 | 1.93 | 3840 | 1495 | 699 | 46.76 |
| M03 | 23 | M | — | 3614.97 | 1.98 | 1920 | 1358 | 393 | 28.94 |
| M04 | 73 | M | CHD | 2442.44 | 1.98 | 1920 | 692 | 291 | 42.05 |
| M05 | 53 | M | CHD | 2419.78 | 1.82 | 1440 | 1013 | 251 | 24.78 |
| M06 | 53 | M | CHD | 3076.19 | 1.75 | 1920 | 733 | 290 | 39.56 |
| M07 | 80 | M | CHD | 3048.77 | 1.90 | 1920 | 1210 | 360 | 29.75 |
| M08 | 14 | M | ASD | 2579.43 | 1.72 | 1920 | 1252 | 231 | 18.45 |
| M09 | 0.8 | M | PDA | 2724.21 | 1.51 | 1920 | 1537 | 249 | 16.20 |
| M10 | 71 | M | MSR | 3094.05 | 1.90 | 1920 | 1411 | 251 | 17.79 |
| F01 | 62 | F | MVPBD | 3421.72 | 1.85 | 1920 | 1432 | 433 | 30.24 |
| F02 | 3 | F | VSD | 3075.96 | 1.85 | 1920 | 1444 | 216 | 14.96 |
| F03 | 43 | F | — | 3190.26 | 1.72 | 1920 | 1510 | 306 | 20.26 |
| F04 | 36 | F | — | 2849.04 | 1.57 | 1920 | 1496 | 311 | 20.79 |

| | | | | | | | | | |
|---|---|---|---|---|---|---|---|---|---|
| F05 | 13 | F | — | 3171.37 | 1.86 | 1920 | 1172 | 308 | 26.28 |
| F06 | 44 | F | — | 3002.40 | 1.71 | 1920 | 1158 | 311 | 26.86 |

[1] CHD: Coronary Heart Disease, ASD: Atrial Septal Defect, PDA: Patent Ductus Arteriosus, MSR: Severe Mitral Regurgitation, MVPBD: Mitral Valve Percutaneous Balloon Dilation, VSD: Ventricular Septal Defect

[2] Median read number per cell

[3] QF: Quality filtering

[4] Copy number alteration size ≥ 2 Mb

**Table S2. List of primers used for library construction**

| Primer ID | Sequence |
|---|---|
| | **i5 Index Primer** |
| **N601** | AATGATACGGCGACCACCGAGATCTACACCTCTCTATTCGTCGGCAGCGTC |
| **N602** | AATGATACGGCGACCACCGAGATCTACACTATCCTCTTCGTCGGCAGCGTC |
| **N603** | AATGATACGGCGACCACCGAGATCTACACGTAAGGAGTCGTCGGCAGCGTC |
| **N604** | AATGATACGGCGACCACCGAGATCTACACACTGCATATCGTCGGCAGCGTC |
| **N605** | AATGATACGGCGACCACCGAGATCTACACAAGGAGTATCGTCGGCAGCGTC |
| **N606** | AATGATACGGCGACCACCGAGATCTACACCTAAGCCTTCGTCGGCAGCGTC |
| **N607** | AATGATACGGCGACCACCGAGATCTACACCGTCTAATTCGTCGGCAGCGTC |
| **N608** | AATGATACGGCGACCACCGAGATCTACACTCTCTCCGTCGTCGGCAGCGTC |
| **N609** | AATGATACGGCGACCACCGAGATCTACACTCGACTAGTCGTCGGCAGCGTC |
| **N610** | AATGATACGGCGACCACCGAGATCTACACTTCTAGCTTCGTCGGCAGCGTC |
| **N611** | AATGATACGGCGACCACCGAGATCTACACCCTAGAGTTCGTCGGCAGCGTC |
| **N612** | AATGATACGGCGACCACCGAGATCTACACGCGTAAGATCGTCGGCAGCGTC |
| **N613** | AATGATACGGCGACCACCGAGATCTACACCTATTAAGTCGTCGGCAGCGTC |
| **N614** | AATGATACGGCGACCACCGAGATCTACACAAGGCTATTCGTCGGCAGCGTC |
| **N615** | AATGATACGGCGACCACCGAGATCTACACGAGCCTTATCGTCGGCAGCGTC |
| **N616** | AATGATACGGCGACCACCGAGATCTACACTTATGCGATCGTCGGCAGCGTC |
| **N801’** | AATGATACGGCGACCACCGAGATCTACACTAAGGCGATCGTCGGCAGCGTC |
| **N802’** | AATGATACGGCGACCACCGAGATCTACACCGTACTAGTCGTCGGCAGCGTC |
| **N803’** | AATGATACGGCGACCACCGAGATCTACACAGGCAGAATCGTCGGCAGCGTC |
| **N804’** | AATGATACGGCGACCACCGAGATCTACACTCCTGAGCTCGTCGGCAGCGTC |
| **N805’** | AATGATACGGCGACCACCGAGATCTACACGGACTCCTTCGTCGGCAGCGTC |
| **N806’** | AATGATACGGCGACCACCGAGATCTACACTAGGCATGTCGTCGGCAGCGTC |
| **N807’** | AATGATACGGCGACCACCGAGATCTACACCTCTCTACTCGTCGGCAGCGTC |
| **N808’** | AATGATACGGCGACCACCGAGATCTACACCGAGGCTGTCGTCGGCAGCGTC |
| **N809’** | AATGATACGGCGACCACCGAGATCTACACAAGAGGCATCGTCGGCAGCGTC |
| **N810’** | AATGATACGGCGACCACCGAGATCTACACGTAGAGGATCGTCGGCAGCGTC |
| **N811’** | AATGATACGGCGACCACCGAGATCTACACGCTCATGATCGTCGGCAGCGTC |
| **N812’** | AATGATACGGCGACCACCGAGATCTACACATCTCAGGTCGTCGGCAGCGTC |
| **N813’** | AATGATACGGCGACCACCGAGATCTACACACTCGCTATCGTCGGCAGCGTC |
| **N814’** | AATGATACGGCGACCACCGAGATCTACACGGAGCTACTCGTCGGCAGCGTC |
| **N815’** | AATGATACGGCGACCACCGAGATCTACACGCGTAGTATCGTCGGCAGCGTC |
| **N816’** | AATGATACGGCGACCACCGAGATCTACACCGGAGCCTTCGTCGGCAGCGTC |
| **N817’** | AATGATACGGCGACCACCGAGATCTACACTACGCTGCTCGTCGGCAGCGTC |
| **N818’** | AATGATACGGCGACCACCGAGATCTACACATGCGCAGTCGTCGGCAGCGTC |
| **N819’** | AATGATACGGCGACCACCGAGATCTACACTAGCGCTCTCGTCGGCAGCGTC |
| **N820’** | AATGATACGGCGACCACCGAGATCTACACACTGAGCGTCGTCGGCAGCGTC |
| **N821’** | AATGATACGGCGACCACCGAGATCTACACCCTAAGACTCGTCGGCAGCGTC |

| | |
|---|---|
| **N822’** | AATGATACGGCGACCACCGAGATCTACACCGATCAGTTCGTCGGCAGCGTC |
| **N823’** | AATGATACGGCGACCACCGAGATCTACACTGCAGCTATCGTCGGCAGCGTC |
| **N824’** | AATGATACGGCGACCACCGAGATCTACACTCGACGTCTCGTCGGCAGCGTC |
| **i7 Index Primer** | |
| **N801** | CAAGCAGAAGACGGCATACGAGATTCGCCTTAGTCTCGTGGGCTCGG |
| **N802** | CAAGCAGAAGACGGCATACGAGATCTAGTACGGTCTCGTGGGCTCGG |
| **N803** | CAAGCAGAAGACGGCATACGAGATTTCTGCCTGTCTCGTGGGCTCGG |
| **N804** | CAAGCAGAAGACGGCATACGAGATGCTCAGGAGTCTCGTGGGCTCGG |
| **N805** | CAAGCAGAAGACGGCATACGAGATAGGAGTCCGTCTCGTGGGCTCGG |
| **N806** | CAAGCAGAAGACGGCATACGAGATCATGCCTAGTCTCGTGGGCTCGG |
| **N807** | CAAGCAGAAGACGGCATACGAGATGTAGAGAGGTCTCGTGGGCTCGG |
| **N808** | CAAGCAGAAGACGGCATACGAGATCAGCCTCGGTCTCGTGGGCTCGG |
| **N809** | CAAGCAGAAGACGGCATACGAGATTGCCTCTTGTCTCGTGGGCTCGG |
| **N810** | CAAGCAGAAGACGGCATACGAGATTCCTCTACGTCTCGTGGGCTCGG |
| **N811** | CAAGCAGAAGACGGCATACGAGATTCATGAGCGTCTCGTGGGCTCGG |
| **N812** | CAAGCAGAAGACGGCATACGAGATCCTGAGATGTCTCGTGGGCTCGG |
| **N813** | CAAGCAGAAGACGGCATACGAGATTAGCGAGTGTCTCGTGGGCTCGG |
| **N814** | CAAGCAGAAGACGGCATACGAGATGTAGCTCCGTCTCGTGGGCTCGG |
| **N815** | CAAGCAGAAGACGGCATACGAGATTACTACGCGTCTCGTGGGCTCGG |
| **N816** | CAAGCAGAAGACGGCATACGAGATAGGCTCCGGTCTCGTGGGCTCGG |
| **N817** | CAAGCAGAAGACGGCATACGAGATGCAGCGTAGTCTCGTGGGCTCGG |
| **N818** | CAAGCAGAAGACGGCATACGAGATCTGCGCATGTCTCGTGGGCTCGG |
| **N819** | CAAGCAGAAGACGGCATACGAGATGAGCGCTAGTCTCGTGGGCTCGG |
| **N820** | CAAGCAGAAGACGGCATACGAGATCGCTCAGTGTCTCGTGGGCTCGG |
| **N821** | CAAGCAGAAGACGGCATACGAGATGTCTTAGGGTCTCGTGGGCTCGG |
| **N822** | CAAGCAGAAGACGGCATACGAGATACTGATCGGTCTCGTGGGCTCGG |
| **N823** | CAAGCAGAAGACGGCATACGAGATTAGCTGCAGTCTCGTGGGCTCGG |
| **N824** | CAAGCAGAAGACGGCATACGAGATGACGTCGAGTCTCGTGGGCTCGG |
| **N601’** | CAAGCAGAAGACGGCATACGAGATATAGAGAGGTCTCGTGGGCTCGG |
| **N602’** | CAAGCAGAAGACGGCATACGAGATAGAGGATAGTCTCGTGGGCTCGG |
| **N603’** | CAAGCAGAAGACGGCATACGAGATCTCCTTACGTCTCGTGGGCTCGG |
| **N604’** | CAAGCAGAAGACGGCATACGAGATTATGCAGTGTCTCGTGGGCTCGG |
| **N605’** | CAAGCAGAAGACGGCATACGAGATTACTCCTTGTCTCGTGGGCTCGG |
| **N606’** | CAAGCAGAAGACGGCATACGAGATAGGCTTAGGTCTCGTGGGCTCGG |
| **N607’** | CAAGCAGAAGACGGCATACGAGATATTAGACGGTCTCGTGGGCTCGG |
| **N608’** | CAAGCAGAAGACGGCATACGAGATCGGAGAGAGTCTCGTGGGCTCGG |
| **N609’** | CAAGCAGAAGACGGCATACGAGATCTAGTCGAGTCTCGTGGGCTCGG |
| **N610’** | CAAGCAGAAGACGGCATACGAGATAGCTAGAAGTCTCGTGGGCTCGG |
| **N611’** | CAAGCAGAAGACGGCATACGAGATACTCTAGGGTCTCGTGGGCTCGG |
| **N612’** | CAAGCAGAAGACGGCATACGAGATTCTTACGCGTCTCGTGGGCTCGG |
| **N613’** | CAAGCAGAAGACGGCATACGAGATCTTAATAGGTCTCGTGGGCTCGG |

| | |
|---|---|
| **N614’** | CAAGCAGAAGACGGCATACGAGATATAGCCTTGTCTCGTGGGCTCGG |
| **N615’** | CAAGCAGAAGACGGCATACGAGATTAAGGCTCGTCTCGTGGGCTCGG |
| **N616’** | CAAGCAGAAGACGGCATACGAGATTCGCATAAGTCTCGTGGGCTCGG |

**Table S3. Fluorescence in situ hybridization results of blood samples showing chromosome loss/gain**

| Donor ID | Chromosome | Type | Abnormal Cell | Total Cells | Ratio (%) |
|---|---|---|---|---|---|
| M01 | 3 | Loss | 58 | 1000 | 5.8 |
| | 3 | Gain | 0 | 1000 | 0 |
| | 8 | Loss | 28 | 1500 | 1.9 |
| | 8 | Gain | 4 | 1500 | 0.27 |
| | 13 | Loss | 14 | 1500 | 0.93 |
| | 13 | Gain | 0 | 1500 | 0 |
| | 18 | Loss | 28 | 1000 | 2.8 |
| | 18 | Gain | 4 | 1000 | 0.4 |
| | 21 | Loss | 35 | 1500 | 2.3 |
| | 21 | Gain | 7 | 1500 | 0.47 |
| M02 | 3 | Loss | 5 | 1000 | 0.5 |
| | 3 | Gain | 14 | 1000 | 1.4 |
| | 8 | Loss | 31 | 1500 | 2.1 |
| | 8 | Gain | 20 | 1500 | 1.3 |
| | 13 | Loss | 38 | 1500 | 2.5 |
| | 13 | Gain | 0 | 1500 | 0 |
| | 18 | Loss | 18 | 1000 | 1.8 |
| | 18 | Gain | 1 | 1000 | 0.1 |
| | 21 | Loss | 52 | 1500 | 3.5 |
| | 21 | Gain | 6 | 1500 | 0.4 |
| F03 | X | Loss | 15 | 1000 | 1.5 |
| | X | Gain | 0 | 1000 | 0 |

**Table S4. Size distributions of unique copy number alterations (CNA) in cell line GM12878**

| CNA Size | 0.6 Mb–2 Mb | 2 Mb–10 Mb | 10 Mb–73.4 Mb | Total |
|---|---|---|---|---|
| **Count** | 35 | 32 | 11 | 78 |
| **Ratio** | 45% | 41% | 14% | 100% |

**Table S5. Copy number alteration (CNA) statistics using simulated data**

| σ | Batch | False Positive (FP) CNA | False Positive Rate | Average Length of FP CNAs (Mb) |
|---|---|---|---|---|
| **0.4** | 1 | 1 | 0.1% | 0.6 |
| | 2 | 2 | 0.2% | 3.1 |
| | 3 | 0 | 0.0% | 0 |
| **0.45** | 1 | 1 | 0.1% | 1 |
| | 2 | 1 | 0.1% | 0.6 |
| | 3 | 1 | 0.1% | 2.0 |
| **0.5** | 1 | 2 | 0.2% | 2.4 |
| | 2 | 1 | 0.1% | 2.2 |
| | 3 | 1 | 0.1% | 3.2 |
| **0.6** | 1 | 2 | 0.2% | 1.3 |
| | 2 | 2 | 0.2% | 2.5 |
| | 3 | 2 | 0.2% | 2.4 |
| **0.7** | 1 | 2 | 0.1% | 2.1 |
| | 2 | 16 | 1.6% | 2.5 |
| | 3 | 5 | 0.5% | 2.3 |
| **0.8** | 1 | 5 | 0.5% | 2.5 |
| | 2 | 15 | 1.5% | 2.2 |
| | 3 | 13 | 1.3% | 2.4 |
